# Unsupervised Discovery of Ancestry Informative Markers and Genetic Admixture Proportions in Biobank-Scale Data Sets

**DOI:** 10.1101/2022.10.22.513294

**Authors:** Seyoon Ko, Benjamin B. Chu, Daniel Peterson, Chidera Okenwa, Jeanette C. Papp, David H. Alexander, Eric M. Sobel, Hua Zhou, Kenneth L. Lange

## Abstract

Admixture estimation plays a crucial role in ancestry inference and genomewide association studies (GWAS). Computer programs such as ADMIXTURE and STRUCTURE are commonly employed to estimate the admixture proportions of sample individuals. However, these programs can be overwhelmed by the computational burdens imposed by the 10^5^ to 10^6^ samples and millions of markers commonly found in modern biobanks. An attractive strategy is to run these programs on a set of ancestry informative SNP markers (AIMs) that exhibit substantially different frequencies across populations. Unfortunately, existing methods for identifying AIMs require knowing ancestry labels for a subset of the sample. This supervised learning approach creates a chicken and the egg scenario. In this paper, we present an unsupervised, scalable framework that seamlessly carries out AIM selection and likelihood-based estimation of admixture proportions. Our simulated and real data examples show that this approach is scalable to modern biobank data sets. Our implementation of the method is called OpenADMIXTURE.

## 1 Introduction

Discovery of ancestral groups by genetic means is of inherent interest for both private genealogies and public population histories.^1^ In addition, genetic ancestry adjustment is a necessary prelude for genome-wide association studies (GWAS) ^2^ seeking DNA sites that influence medical or anthropomorphic traits. Without this safeguard, population stratification can lead to a false association between a trait and a SNP (single nucleotide polymorphism) predictor.^3,4,5^ Ancestry adjustment can be handled by adding a few principal components (PCs) of the SNP genotype matrix as trait predictors. Alternatively, one can substitute admixture proportions (coefficients) in place of PCs. Because admixture coefficients are the proportions of an individual’s ancestry from each of several founding populations, they are usually more interpretable than PCs.

Estimation of ancestry coefficients is carried out simultaneously with maximum likelihood estimation of allele frequencies when the underlying populations are latent rather than explicit. ADMIXTURE ^6^ is the most widely used likelihood-based software. ADMIXTURE directly maximizes the likelihood of the genotype matrix via alternating sequential quadratic programming with quasi-Newton acceleration^7^. Another popular package, STRUCTURE^8^, implements Bayesian inference; recent extensions of the Bayesian approach include fast-Structure^9^ and TeraStructure^10^. The EIGENSTRAT software^2^ is the primary vehicle for PC extraction from the genotype matrix. One can also achieve dimensionality reduction by approximating a low-rank linear subspace of the row space of the admixture proportion matrix and then performing matrix decomposition via alternating least squares software^11^. The ALStructure software implementing matrix decomposition can be further accelerated by invoking randomized linear algebra and routines specifically designed for accessing compressed versions of the genotype matrix in the SCOPE software^12^.

As the size of genomic data grows, these methods suffer. In particular, most of the methods fail on the UK Biobank data^13^, which contains {0, 1, 2} genotypes on about half a million British individuals across millions of SNPs. The genotype matrix in PLINK format alone requires around 70 GB of storage. The only known software that is capable of handling these data is SCOPE^12^, which avoids holding large intermediate matrices in memory. However, SCOPE’s preprocessing of the genotype matrix to speed up matrix multiplication still requires at least 250 GB of memory (RAM).

One can make ancestry estimation more efficient by limiting analysis to ancestry informative markers (AIMs)^14,15^. Early AIM sets included tens to hundreds of AIMs^16,17,18,19^. Even at this crude scale, it is possible to recover admixture coefficients that correlate well (74-92%) with those delivered by the full set of SNPs^20^. AIM based methods exploit F statistics, absolute allele frequency differences, principal component loadings, and informativeness in ancestry assignment^21,22^. To their disadvantage, most AIM selection methods are supervised and based on self-reported labels. In biobank-scale data, such labels should be viewed with suspicion.

In the current paper we advocate first selecting AIMs in an unsupervised way through a sparse *K*-means clustering algorithm ^23^. We will refer to this algorithm by the acronym SKFR (sparse *K*-means with feature ranking). SKFR performs hard clustering and feature selection jointly and is scalable to biobank data. Given the selected AIMs, we run a new version of ADMIXTURE^6^ that leverages the computational advances of the Julia programming language^24^. We call this new package OpenADMIXTURE^25^, due in part to its open-source status. OpenADMIXTURE incorporates both SKFR and admixture estimation, supports multithreading, and acts directly on the input genotype matrix. The maximum memory demand is less than 120% the size of the input genotype file. For example, our analysis of the UK Biobank data with 500,000 individuals and 600,000 SNPs requires only 73 GB of memory, versus the 250 GB required by SCOPE. OpenADMIXTURE also supports graphics processing unit (GPU) acceleration. Runtimes and results of OpenADMIXTURE are comparable to those of SCOPE but within the RAM limitations of more typical computers. Furthermore, OpenADMIXTURE retains the advantages of a likelihood-based analysis. SKFR is generally useful in feature selection across a wide variety of clustering applications beyond genetics. An independent Julia-based package to efficiently perform SKFR on general data sets is available^26^.

## 2 Methods

### 2.1 Sparse K-Means with Feature Ranking (SKFR)

SKFR selects and ranks a predetermined number of features that drive *K*-means clustering^23^. Feature selection and clustering performance are intertwined. In our case, features are standardized SNP genotypes displayed in an *I × J* matrix ***X*** = (*x*_*ij*_). Rows correspond to samples and columns to features. Standardization of columns to have mean 0 and variance 1 puts all features on the same footing. Given a fixed number of clusters *K*, the goal is to assign each individual *i* and its corresponding row 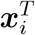 of ***X*** to the cluster *C*_*k*_ minimizing the loss

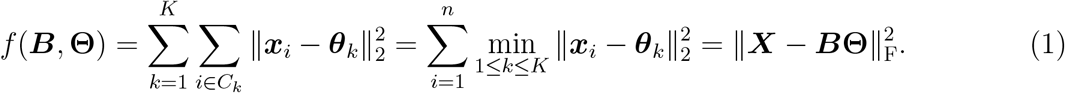

Here the matrix ***B*** *∈* {***M*** *∈* {0, 1}^*I×K*^ : ***M* 1**_*K*_ = **1**_*I*_} conveys cluster membership, the matrix **Θ** *∈* ℝ^*K×J*^ conveys cluster centers, and ∥ *·* ∥_F_ denotes the Frobenius norm. The *k*-th row 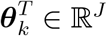 of **Θ** is the center of cluster *C*_*k*_. In SKFR with sparsity level *S*, at most *S* columns of **Θ** are allowed to be nonzero. The SKFR procedure (see Algorithm 1) cycles through the following three steps until convergence: (a) update the cluster centers, (b) rank and select features according to their contribution to the loss, (c) re-assign samples to clusters according to the selected features. In Algorithm 1, the information criterion *h*_*j*_ measures the drop in the loss when feature *j* is designated informative. The clusters are initialized by the *K*-means++ scheme^27^. Initial cluster centers emerge after steps (b) and (c) are performed on the standardized matrix ***X***. Section 2.1.1 sketches modification of the algorithm to handle missing data.

On convergence, the SKFR algorithm yields (a) a ranked list *L* of selected AIMs, (b) hard clustering assignments ***B*** of each sample to one and only one cluster, and (c) cluster centers **Θ**. A new set of PLINK files containing only the selected AIMs is generated via the `SnpArrays` Julia package.

#### Algorithm 1 SKFR Algorithm

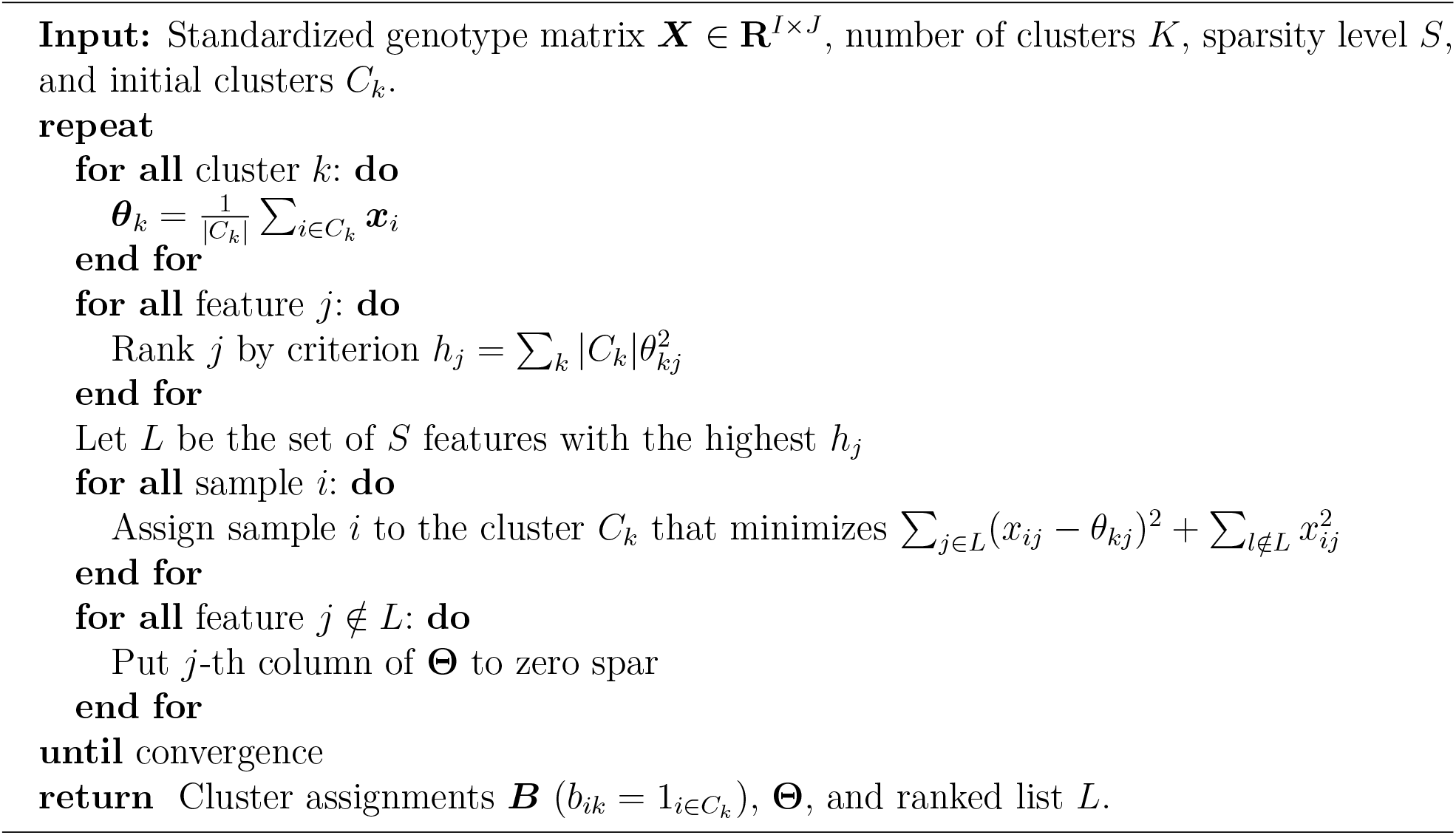

#### 2.1.1 Missing Genotypes

Genotype data often include missing values. In practice, genotype imputation precedes GWAS. However, imputing genotypes at biobank scale is extremely resource and computation intensive, traditionally taking days to months on a cluster. Modern software^28,29^ has reduced this bottleneck. Following Chi et al. ^30^, we extend SKFR to incorporate missing data in a mathematically principled way. Let Ω ⊂ {1, …, *I*} *×* {1, …, *J* } denote the subset of indices corresponding to the observed entries of ***X***. In this notation, the modified loss is

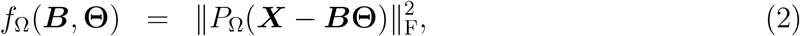

where *P*_Ω_(***M***) zeros all entries of a matrix ***M*** not in Ω.

The quickest route to minimization of the loss passes through the majorization-minimization (MM) principle^31,32,33^. At iteration *n* of a search, we construct the surrogate function 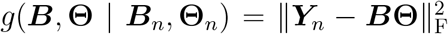 majorizing the loss, where 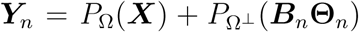 and Ω^⊥^ denotes the set of observed values complementary to the set of missing values. In other words, we leave observed values untouched and impute missing values by their predicted values based on the centers of their current cluster assignments. The MM principle guarantees that minimizing the surrogate reduces the loss. This monotonic algorithm is summarized in Algorithm 2. The current code differs from the code presented in Zhang et al. ^23^ by standardizing the genotype matrix beforehand using observed values rather than repeatedly standardizing on the fly using both observed and imputed values. The current implementation is more efficient and mathematically sound.

##### Algorithm 2 SKFR Algorithm Incorporating Missing Genotypes

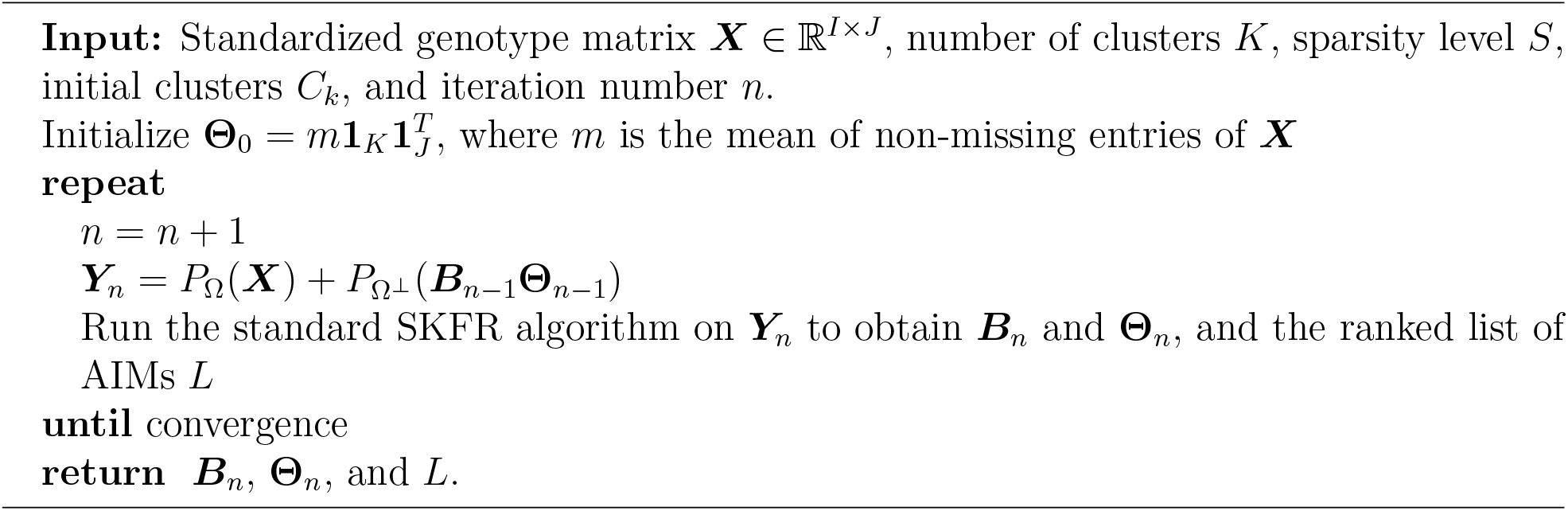

### 2.2 Estimation of Admixture Proportions

In contrast to hard clustering, soft clustering estimates the probability of a sample belonging to each of the *K* clusters. Soft clustering algorithms like ADMIXTURE^6^ better account for ambiguities than hard clustering and in GWAS more realistically adjust for population structure. In this section we describe a Julia implementation of ADMIXTURE that capitalizes on parallel processing and GPU support. Recall that ADMIXTURE simultaneously estimates a population-specific allele frequency matrix ***P*** *∈* ℝ^*K×J*^ and an individual-specific admixture matrix ***Q*** *∈* ℝ^*K×I*^ by maximizing the loglikelihood

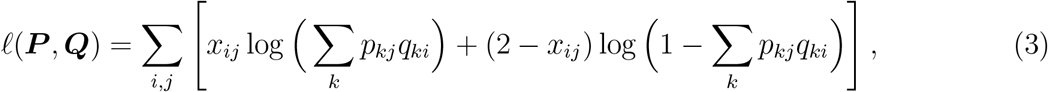

Here each raw genotype *x*_*ij*_ follows a Binomial(2, *Σ* _*k*_ *p*_*kj*_*q*_*ki*_) distribution, where the parameters satisfy the constraints 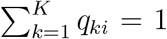 and 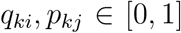. Maximization is carried out by block ascent, alternating updates of ***P*** and ***Q*** by sequential quadratic programming with quasi-Newton acceleration^7^.

Given an objective function *f* (***x***), sequential quadratic programming finds the next iterate ***x***_*n*+1_ = ***x***_*n*_ + **Δ** by minimizing the second-order approximation

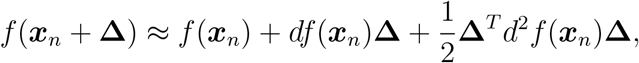

with respect to **Δ** subject to relevant constraints. Here *df* (***x***) is the first differential (transposed gradient) and *d*^2^*f* (***x***) is the second differential (Hessian) of *f* (***x***). With linear constraints and upper and lower bounds, one can exploit a standard pivoting strategy to solve this quadratic program^34^. ADMIXTURE accomplishes precisely this with the objective equal to the negative loglikelihood. In the block ascent updates, the SNP-specific allele frequencies and individual-specific admixture proportions are parameters that can be separated. This results in an overall computational complexity of *O*[(*I* + *J*)*K*^3^] for the quadratic programs, which is negligible compared to the bottleneck of *O*(*IJK*^2^) in computing Hessians, as *K* ≪ *I, J*. Admixture proportions are initialized by five iterations of FRAPPE’s EM (expectation-maximization) algorithm ^35^.

OpenADMIXTURE leverages Julia to achieve higher performance. The core computations of sequential quadratic programming now exploit tiling to maximize locality and avoid cache misses. Users can choose to offload most computations to graphics processing units (GPUs) for further speedup. OpenADMIXTURE’s default setting declares convergence when the relative change in loglikelihoods is less than 10^−7^. Supervised inference is possible by fixing ***P*** and updating only ***Q*** or fixing ***Q*** and updating only ***P***. The former is pertinent when admixture proportions are sought and predefined allele frequencies from reference populations are used. The latter is pertinent when allele frequencies are sought for the reference populations whose admixture coefficients are fixed.

### 2.3 Software Input and Output

Our OpenADMIXTURE package internally runs a pipeline of SKFR and then admixture estimation. As mentioned above, SKFR is also available as a standalone package for use on general data sets^26^. It is also possible to run OpenADMIXTURE using all available SNPs and bypass AIM discovery through SKFR. Given these considerations the input and output conventions adopted by OpenADMIXTURE are the following.

#### For SKFR

**Input**: A single set (`bed`, `fam`, `bim`) or collection of PLINK binary files, the number of clusters, and the number of AIMs.

**Output**: A single set of PLINK binary files containing only the selected AIMs, and a file containing hard clustering results, with each row indicating the cluster to which a sample is assigned. A filtered PLINK file containing only the AIMs is optional.

#### For Admixture Estimation

**Input**: A single set (`bed`, `fam`, `bim`) or collection of PLINK binary files, possibly filtered to contain only the selected AIMs under SKFR, and the number of clusters.

**Output**: A `P` file with each row recording the cluster-specific allele frequencies of an AIM, and a `Q` file with each row recording the estimated admixture proportions of an individual.

### 2.4 Computational Tactics

#### 2.4.1 Direct Computation on the PLINK BED File

The PLINK BED format^36^ stores SNP genotypes in a compressed binary format, with 00, 10, 11, 01 representing the 0, 1, 2, and missing genotypes, respectively. Our software exploits this structure to reduce memory usage through the OpenMendel^37^ package `SnpArrays`. A special data structure is designed so that even if data files are separated by chromosome, the whole is treated as a single array through a high-level interface.

For SKFR, imputation and standardization are lazily evaluated using the compressed PLINK genotypes, the means and standard deviations of each SNP, and stored values of the cluster centers, updated on each iteration of Algorithm 2. SIMD (single instruction, multiple data)-vectorization and multithreading are supported for computing distances to cluster centers. Since the genotype matrix is memory-mapped through `SnpArrays`, memory usage is minimized even with multithreading. Direct computation on compressed genotypes, including all GPU kernels, is also implemented in OpenADMIXTURE. Thus, OpenADMIXTURE can analyze 16-32 times more data per GPU than GPU-based admixture software such as Mantes et al. ^38^, whose GPU kernels require a single precision or double precision genotype matrix as input.

#### 2.4.2 Initialization

Initialization of SKFR follows the *K*-means++^27^ recommendation. For ADMIXTURE, five EM iterations as employed in the FRAPPE software^35^ gives fast, rough estimates. Although one could plausibly use the allele frequencies of the SKFR clusters as initial values, hard clustering sometimes produces small clusters with extreme frequencies. More advanced techniques such as power *K*-means^39^ may well yield even better initialization.

#### 2.4.3 Updates of Cluster Centers in SKFR

After updating cluster labels, we update cluster centers based on only the samples reassigned to new clusters. This is advantageous because just a small proportion of samples changes labels at any iteration. To their detriment, downdating and updating of centers involves all SNPs, not just the currently informative ones. For a sample *i* changing its assignment to *C*_*k*′_ from *C*_*k*_, suppose |*C*_*k*′_ | = *n*_*k*′_ and |*C*_*k*_| = *n*_*k*_ after reassignment. We then update the cluster center 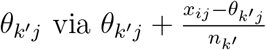 and downdate the cluster center 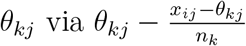.

#### 2.4.4 Acceleration with Missing Genotypes in SKFR

The originally proposed method for imputation of missing genotypes in Zhang et al. ^23^ mandates standardization of ***Y***_*n*_ at each iteration of Algorithm 2. To simplify computation, we now standardize the genotype matrix ***X***_*n*_ only once before iterations commence. We also run just a single inner iteration of SKFR per iteration of *k*-pod imputation for better performance. The descent property is maintained.

#### 2.4.5 Warm Start for a Path of Different Sparsity Levels S

It is often desirable to explore a variety of AIM sparsity levels *S* on the same data set. This can be done efficiently by starting with the highest level *S*_max_ desired and gradually decreasing *S*. The results from a given *S* are then invoked to warm start computations at the next lower level of *S*. OpenADMIXTURE’s ranking of AIMs facilitates this tactic.

#### 2.4.6 Tiling in OpenADMIXTURE

In the CPU version of OpenADMIXTURE software, recursive tiling based on cache-oblivious algorithms^40^ is employed for updating gradients and Hessians of the loglikelihood. These computations are carried out in blocks of small size (for example, smaller than 64 *×* 64) in the *I × J* index space of the Hessian computation. Tiling improves the locality of execution, thus increasing cache efficiency and consequently program speed.

#### 2.4.7 GPU Acceleration of OpenADMIXTURE

For OpenADMIXTURE, GPU kernels are designed to harness the Julia `CUDA` package^41^ in computing the loglikelihood and its gradient and Hessian. The gradient and Hessian can be computed in single precision to speed up computations.

#### 2.4.8 Multithreading and Automatic Vectorization

Multithreading is implemented using native commands supported in Julia in conjunction with the automatic vectorization package, `LoopVectorization`, which makes effective use of vector instructions such as AVX-512 on Intel CPUs through the macro @turbo in Julia.

### 2.5 Postprocessing for Performance Evaluation

#### 2.5.1 Permutation Matching of Clusters

Clustering results derived from two separate algorithms can be compared by various statistics. Any such statistic should be invariant under permutation of cluster labels and match similar clusters. We carry out matching following the approach of Behr et al. ^42^. The similarity between cluster *m* of ***Q***^1^ and cluster *n* of ***Q***^2^ is quantified by

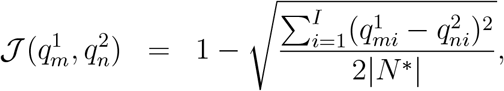

where 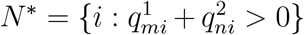. Cluster matching can be formulated as an assignment problem maximizing the criterion 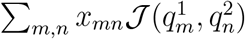 subject to the constraints *x*_*mn*_ *∈* {0, 1} and 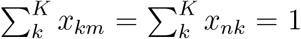. In practice, the domain of *x*_*mn*_ is relaxed to the unit interval, and the problem is solved by linear programming via JuMP^43^, Julia’s mathematical optimization package.

#### 2.5.2 Visualization

We visualize estimates for admixture proportions as stacked bar plots. The clusters in each run are matched for easy comparison. To determine the order of samples, we rely on hierarchical clustering with complete linkage based on the OpenADMIXTURE ***Q*** estimates. The samples are ordered within each population, and the populations are ordered based on hierarchical clustering of cluster centers. The same is done for superpopulations whenever applicable.

## 3 Analyses of Large-Scale Real Data Sets

We ran OpenADMIXTURE, encompassing both SKFR and admixture estimation, on four independent data sets: 1000 Genomes Project (TGP)^44,45^, Human Genome Diversity Project (HGDP)^46,47^, Human Origins (HO)^48^, and UK Biobank (UKB). The TGP data set consists of the 2012-01-31 Omni Platform genotypes confined to unrelated individuals with at least a 95% genotyping success rate and SNPs with at least a 1% minor allele frequency (MAF). The filtered data set contains 1718 individuals and 1,854,622 SNPs. The original VCF-formatted data are converted to PLINK BED format. Samples are labeled as belonging to one of 26 populations, which are grouped into five superpopulations designated African, Admixed American, East Asian, European, and South Asian. The HGDP data set contains the individuals in the Stanford H952 data set with greater than a 95% genotyping success rate and SNPs with at least a 1% MAF. The HGDP data contain 642,951 SNPs and 940 individuals across 32 populations, which are grouped into seven continental superpopulations: Europe, Middle East, Central South Asia, East Asia, Africa, America, and Oceania. The HO data set is filtered to include only samples with at least a 99% genotype success rate and SNPs with at least a 5% MAF. The HD data contain 385,089 SNPs for 1931 people across 163 populations. Continental population labels are not provided. For the UK Biobank (UKB) data set^13^, we filtered bulk genotypes to include individuals with at least a 95% genotyping success rate and SNPs with at least a 1% MAF. The resulting data include 488,154 individuals and 610,741 SNPs.

### 3.1 Evaluation of Results

#### 3.1.1 Hard Clustering

To evaluate the clustering performance of SKFR, we ran it on the TGP data set with *K* = 8, HGDP with *K* = 10, and HO with *K* = 14 clusters, choices consistent with previous analyses of these data^10,11,12^. Recall that within the data set each individual is labeled as coming from one of 26 populations and one of five superpopulations. We tried 10 different initializations for sparsity level *S* = 100, 000 and chose the best clustering according to the loss function of equation (2). Then we successively decremented *S* to 80,000, 60,000, 40,000, 20,000, 10,000, and 5000 using the warm start tactic described in Section 2.4.5. We computed the adjusted Rand index (ARI)^49,50^ and the normalized mutual information (NMI)^51^ between our hard clusterings and the five superpopulation labels originally assigned within the data set to the samples. Although these two metrics are rather opaque, they do allow the number of clusters to differ in each clustering. These measures were also computed for the baseline *K*-means clustering with all SNPs included. The baseline results also reflect 10 different initializations.

Table 1 for the TGP data shows that SKFR’s hard clusterings clearly outperform the baseline *K*-means results and that the SKFR results are stable across a wide range of selected SNPs. When we exclude the admixed American (AMR) superpopulation in our assessment, our clusters perfectly capture the four remaining superpopulations. It appears that including uninformative SNPs or admixed samples creates unwanted noise that obscures true clusters. With HGDP and HO data sets (Tables 2 and 3, respectively), ARI and NMI of SKFR is comparable to but slightly worse than those of the baseline *K*-means. For the HGDP data this anomalous result may stem from the admixed nature of the HGDP superpopulations. It is also noteworthy that the HO data include 163 different population labels. With HGDP and HO data sets, ARI and NMI decrease as we choose more AIMs, with up to 100,000 AIMs selected. This anomaly may have two sources. First, we are relying on possibly inaccurate self-reported population labels. Second, we are hard labeling individuals who may be admixed. Unlike TGP data where it is straightforward to distinguish admixed population (one continental label is literally “Admixed Americans”), it is much more difficult to isolate less admixed population from continental labels of HGDP, as it intentionally collected samples with more diverse background. In case of HO, we are using 163 labels without any provided continental labels.

**Table 1:** Hard clustering performance on the TGP data with all samples, relative to the five superpopulation labels, and ignoring the admixed American (AMR) samples, relative to the remaining four superpopulation labels. Performance is evaluated using the adjusted Rand index (ARI) and the normalized mutual information (NMI). The best value in each column is in **bold**, except when the maximum value of 1.0 is reached.

| #AIMs | All samples |  | Without AMR |  |
| --- | --- | --- | --- | --- |
|  | ARI | NMI | ARI | NMI |
| 5,000 | 0.824 | 0.839 | 1.000 | 1.000 |
| 10,000 | **0.825** | 0.840 | 1.000 | 1.000 |
| 20,000 | **0.825** | 0.840 | 1.000 | 1.000 |
| 40,000 | **0.825** | **0.841** | 1.000 | 1.000 |
| 60,000 | 0.824 | 0.840 | 1.000 | 1.000 |
| 80,000 | **0.825** | 0.840 | 1.000 | 1.000 |
| 100,000 | 0.822 | 0.837 | 1.000 | 1.000 |
| All SNPs (baseline $K$-means) | 0.575 | 0.726 | 0.726 | 0.802 |

**Table 2:** Hard clustering performance on the HGDP data with all samples, relative to the seven continental labels. Performance is evaluated using the adjusted Rand index (ARI) and the normalized mutual information (NMI). The best value in each column is in **bold**.

| #AIMs | ARI | NMI |
| --- | --- | --- |
| 5,000 | 0.0861 | 0.209 |
| 10,000 | 0.0861 | 0.209 |
| 20,000 | 0.0861 | 0.209 |
| 40,000 | 0.0858 | 0.207 |
| 60,000 | 0.0855 | 0.206 |
| 80,000 | 0.0855 | 0.206 |
| 100,000 | 0.0846 | 0.205 |
| All SNPs (baseline $K$-means) | **0.0898** | **0.216** |

**Table 3:** Hard clustering performance on the HO data with all samples, relative to the 163 population labels. Performance is evaluated using the adjusted Rand index (ARI) and the normalized mutual information (NMI). The best value in each column is in **bold**.

| #AIMs | ARI | NMI |
| --- | --- | --- |
| 5,000 | 0.111 | 0.581 |
| 10,000 | 0.111 | 0.580 |
| 20,000 | 0.111 | 0.579 |
| 40,000 | 0.109 | 0.575 |
| 60,000 | 0.098 | 0.565 |
| 80,000 | 0.083 | 0.548 |
| 100,000 | 0.074 | 0.539 |
| All SNPs (baseline $K$-means) | **0.131** | **0.615** |

#### 3.1.2 Admixture Estimation

We recorded concordance with ancestry labels included in the data sets as a performance measure for soft clustering under OpenADMIXTURE. We also trained a Softmax (multinomial logistic) classifier to predict superpopulation labels using TGP data with the inferred admixture proportions as predictors. Since the results are continuous proportions rather than hard clusters, cross-entropy is a reasonable measure of error. We additionally matched clusters as discussed in Section 2.5.1 and computed root-mean-square error (RMSE) compared to the OpenADMIXTURE estimates with all SNPs included.

Tables 4-6 display our complex findings for the HGDP, HO, and TGP data sets, respectively. The accuracy of OpenADMIXTURE classification with a limited number of AIMs is roughly comparable to that of SCOPE, which always uses all SNPs. In general, cross-entropy decreases (improves) as we select more AIMs in OpenADMIXTURE’s inference. In particular for the HO and TGP data sets, the OpenADMIXTURE estimates with 60,000 or more AIMs outperforms SCOPE. The RMSEs of SCOPE and AIM-driven OpenADMIXTURE are also roughly comparable within each of the three data sets. SCOPE does somewhat better on the HO data, while OpenADMIXTURE does better on the HGDP and TGP data. Again, we stress that the admixed nature of the data may obscure the value of limiting analysis to AIMs and cloud the choice of the optimal number of AIMs.

**Table 4:** Performance of OpenADMIXTURE estimation on the HGDP data set, as measured by training accuracy, cross-entropy with the seven continental labels, and RMSE from the baseline that includes all SNPs in the admixture estimation. Also, results for SCOPE on the complete HGDP data set. “All SNPs (baseline)” omits SKFR. The best value in each column is in **bold**.

| #AIMs | Accuracy | Cross-Entropy | RMSE from Baseline |
| --- | --- | --- | --- |
| 5,000 | 0.463 | 971 | 0.104 |
| 10,000 | 0.469 | 962 | 0.108 |
| 20,000 | 0.470 | 961 | 0.117 |
| 40,000 | 0.467 | 959 | **0.037** |
| 60,000 | 0.463 | 958 | 0.081 |
| 80,000 | 0.470 | 955 | 0.098 |
| 100,000 | 0.464 | 955 | 0.097 |
| All SNPs (baseline) | 0.467 | **953** | — |
| SCOPE | **0.475** | 962 | 0.039 |

**Table 5:** Performance of OpenADMIXTURE estimation on the HO data set, as measured by training accuracy, cross-entropy with the 163 population labels, and RMSE from the baseline that includes all SNPs in the admixture estimation. Also, results for SCOPE on the complete HO data set. “All SNPs (baseline)” omits SKFR. The best value in each column is in **bold**.

| #AIMs | Accuracy | Cross-Entropy | RMSE from Baseline |
| --- | --- | --- | --- |
| 5,000 | 0.225 | 6610 | 0.121 |
| 10,000 | 0.246 | 6349 | 0.105 |
| 20,000 | 0.291 | 6101 | 0.110 |
| 40,000 | 0.320 | 5897 | 0.101 |
| 60,000 | 0.328 | **5851** | 0.093 |
| 80,000 | **0.330** | 5869 | 0.087 |
| 100,000 | 0.323 | 5975 | 0.102 |
| All SNPs (baseline) | 0.325 | 5951 | — |
| SCOPE | 0.275 | 6486 | **0.077** |

**Table 6:** Performance of OpenADMIXTURE estimation on the TGP data set, as measured by training accuracy, cross-entropy with the five (four without admixed Americans [AMR]) continental labels, and RMSE from the baseline that includes all SNPs in the admixture estimation. Also, results for SCOPE on the complete TGP data set. “All SNPs” is used as baseline and omits SKFR. The best value in each column is in **bold**; accuracy has a maximum value of 1.0.

| #AIMs | All samples |  | Without AMR |  |  |
| --- | --- | --- | --- | --- | --- |
|  | Accuracy | Cross-Entropy | Accuracy | Cross-Entropy | RMSE from Baseline |
| 5,000 | 0.941 | 313 | 1.000 | 37.4 | 0.234 |
| 10,000 | 0.947 | 295 | 0.998 | 34.9 | 0.185 |
| 20,000 | 0.947 | 284 | 1.000 | 33.4 | 0.152 |
| 40,000 | 0.953 | 274 | 1.000 | 31.7 | 0.164 |
| 60,000 | 0.971 | 242 | 1.000 | 29.0 | 0.043 |
| 80,000 | 0.968 | 241 | 1.000 | 30.5 | 0.033 |
| 100,000 | 0.969 | 241 | 1.000 | 30.1 | **0.027** |
| All SNPs | **0.980** | **232** | 1.000 | **27.3** | — |
| SCOPE | 0.979 | 248 | 1.000 | 32.1 | 0.044 |

The TGP data demonstrate the value of excluding samples known to be admixed. Table 6 shows that OpenADMIXTURE classification is perfect for the non-AMR individuals with at least 20,000 AIMs selected. The table also shows better cross-entropy for classifying non-admixed samples versus all samples. Table 7 reinforces these findings by omitting AMR samples during SKFR AIM selection prior to admixture analysis. The table shows slightly better classification performance than the performance recorded in Table 6 with the same number of AIMs.

**Table 7:**
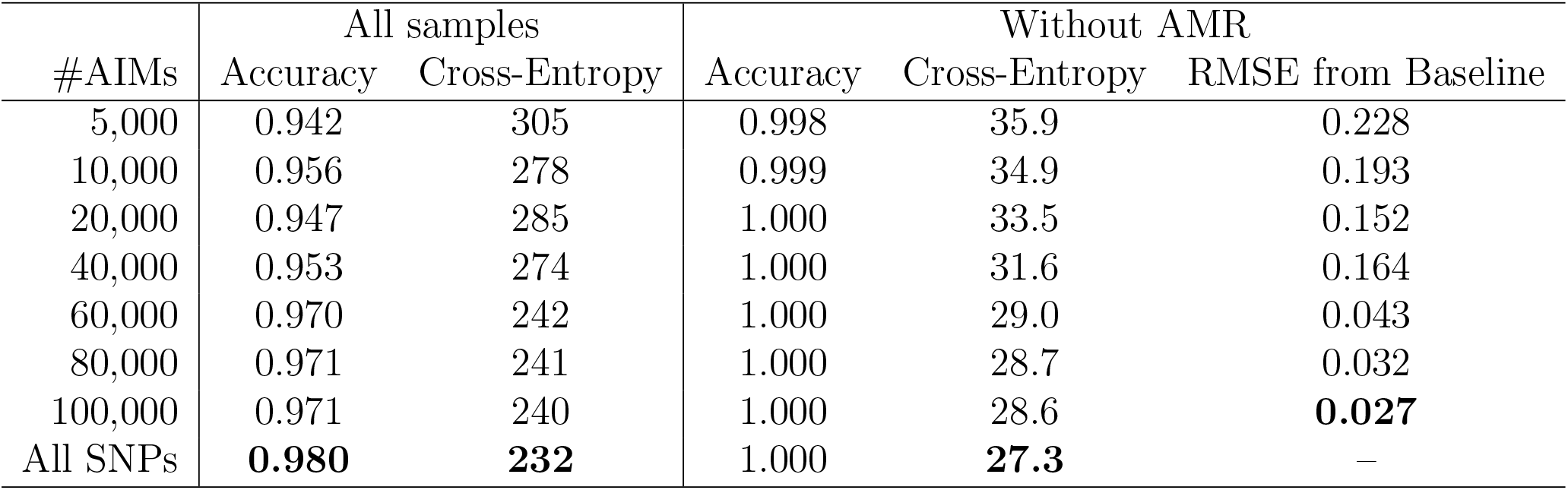
Performance of OpenADMIXTURE on the TGP data set ignoring admixed American samples during the SKFR stage, as measured by training accuracy, cross-entropy with the five (four without admixed Americans [AMR]) continental labels, and RMSE from the baseline that includes all SNPs in the admixture estimation. “All SNPs” is used as baseline and omits SKFR. The best value in each column is in **bold**; accuracy has a maximum value of 1.0.

| #AIMs | All samples |  | Without AMR |  |  |
| --- | --- | --- | --- | --- | --- |
|  | Accuracy | Cross-Entropy | Accuracy | Cross-Entropy | RMSE from Baseline |
| 5,000 | 0.942 | 305 | 0.998 | 35.9 | 0.228 |
| 10,000 | 0.956 | 278 | 0.999 | 34.9 | 0.193 |
| 20,000 | 0.947 | 285 | 1.000 | 33.5 | 0.152 |
| 40,000 | 0.953 | 274 | 1.000 | 31.6 | 0.164 |
| 60,000 | 0.970 | 242 | 1.000 | 29.0 | 0.043 |
| 80,000 | 0.971 | 241 | 1.000 | 28.7 | 0.032 |
| 100,000 | 0.971 | 240 | 1.000 | 28.6 | **0.027** |
| All SNPs | **0.980** | **232** | 1.000 | **27.3** | — |

For the UKB data with *K* = 4 clusters, we computed the accuracy of the Softmax classifier with three sets of labels. The first set (L1) uses all 22 raw labels. The second (L2) uses the eight labels British, Irish, Indian, Pakistani, Bangladeshi, Caribbean, African, and Chinese for roughly homogeneous populations, and removes mixed or uncertain labels such as “mixed” or “other”. The third set (L3) merges L2’s populations into four continental groupings, British-Irish, Indian-Pakistani-Bangladeshi, Caribbean-African, and Chinese. Table 8 reports this classification accuracy for OpenADMIXTURE with *S* = 100, 000 AIMs, and for SCOPE using all SNPs.

**Table 8:** Admixture estimation accuracy of OpenADMIXTURE with 100,000 AIMs and SCOPE on the UKB data set, relative to three labeling sets, L1–L3. L1 uses all individuals along with the population labels assigned by the data set (22 total labels); L2 removes individuals with ambiguous labels such as “other”, and restricts the labels to eight roughly homogeneous populations; and L3 uses four continental labels on the individuals in L2. Higher accuracy is better, with 1.0 the maximum value.

|  | L1 | L2 | L3 |
| --- | --- | --- | --- |
| OpenADMIXTURE (100,000 AIMs) | 0.911 | 0.963 | 0.999 |
| SCOPE | 0.914 | 0.964 | 0.999 |

#### 3.1.3 Visualization

Figures 1-3 depict the inferred admixture proportions for the TGP, HGDP, and HO data sets. Each figure includes three graphs: first, the result from OpenADMIXTURE with all SNPs; second, from OpenADMIXTURE with 100,000 AIMs; and third, from SCOPE.

**Figure 1:**
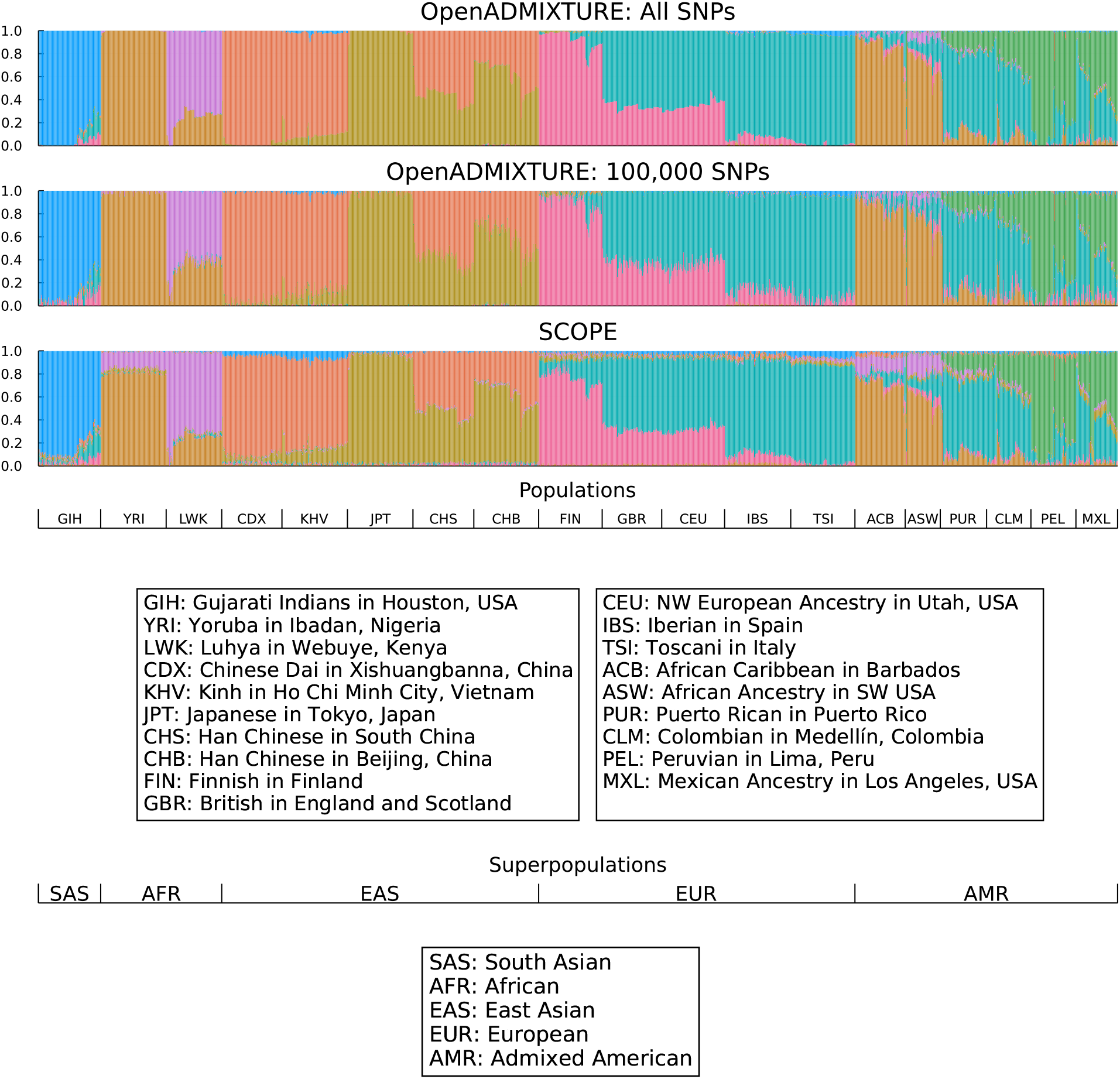
Estimated ancestry of TGP data samples using (a) OpenADMIXTURE with all SNPs, (b) OpenADMIXTURE with 100,000 AIMs, and (c) SCOPE. These are stacked bar plots, with the y-axis indicating the proportion of total ancestry. The x-axis runs over all samples; the population labels originally assigned to these samples within the data set are provided in the lower sections of the figure.

**Figure 2:**
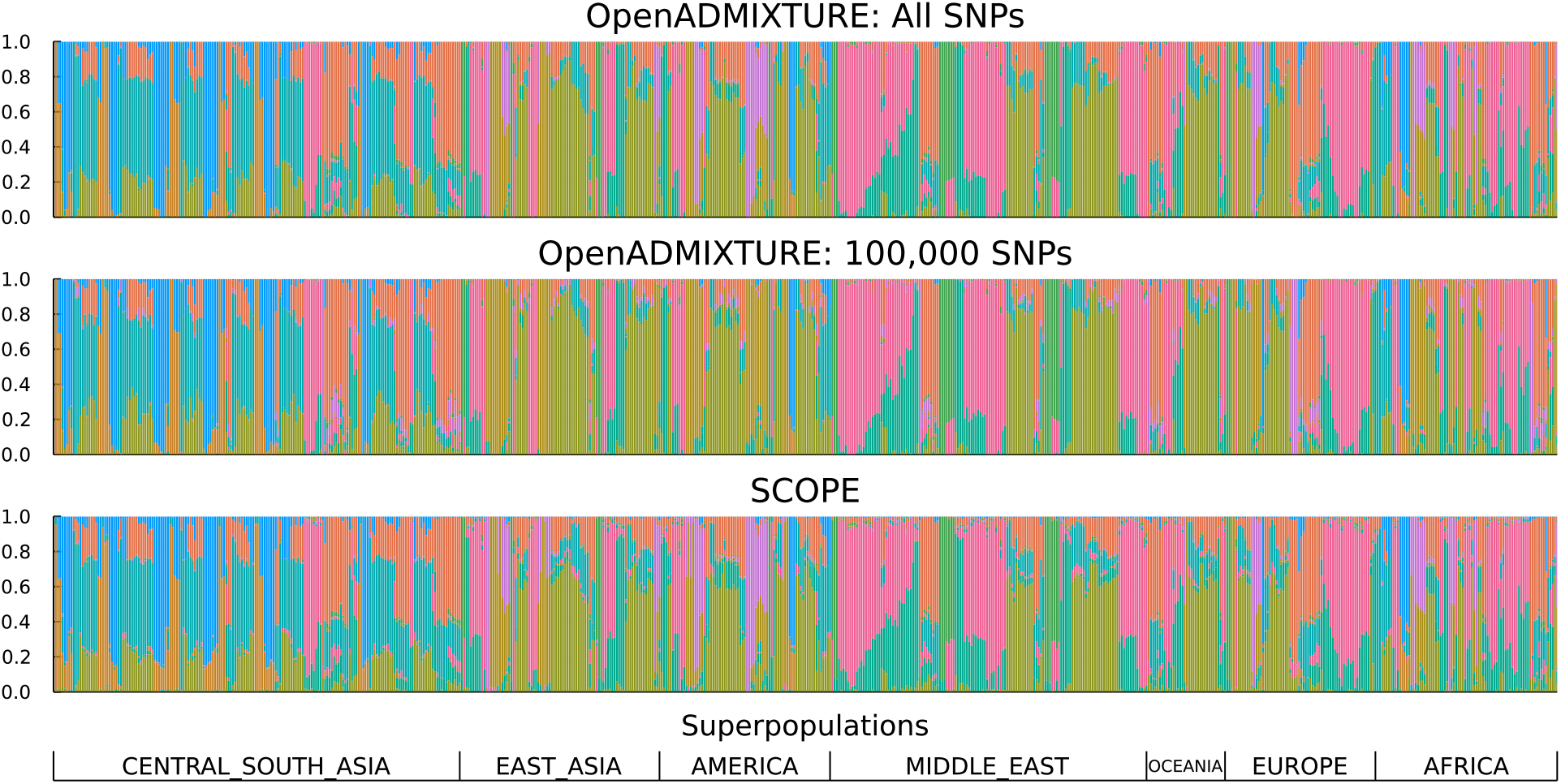
Estimated ancestry of HGDP data samples using (a) OpenADMIXTURE with all SNPs, (b) OpenADMIXTURE with 100,000 AIMs, and (c) SCOPE. These are stacked bar plots, with the y-axis indicating the proportion of total ancestry. The x-axis runs over all samples; the population labels originally assigned to these samples within the data set are provided in the lower section of the figure.

**Figure 3:**
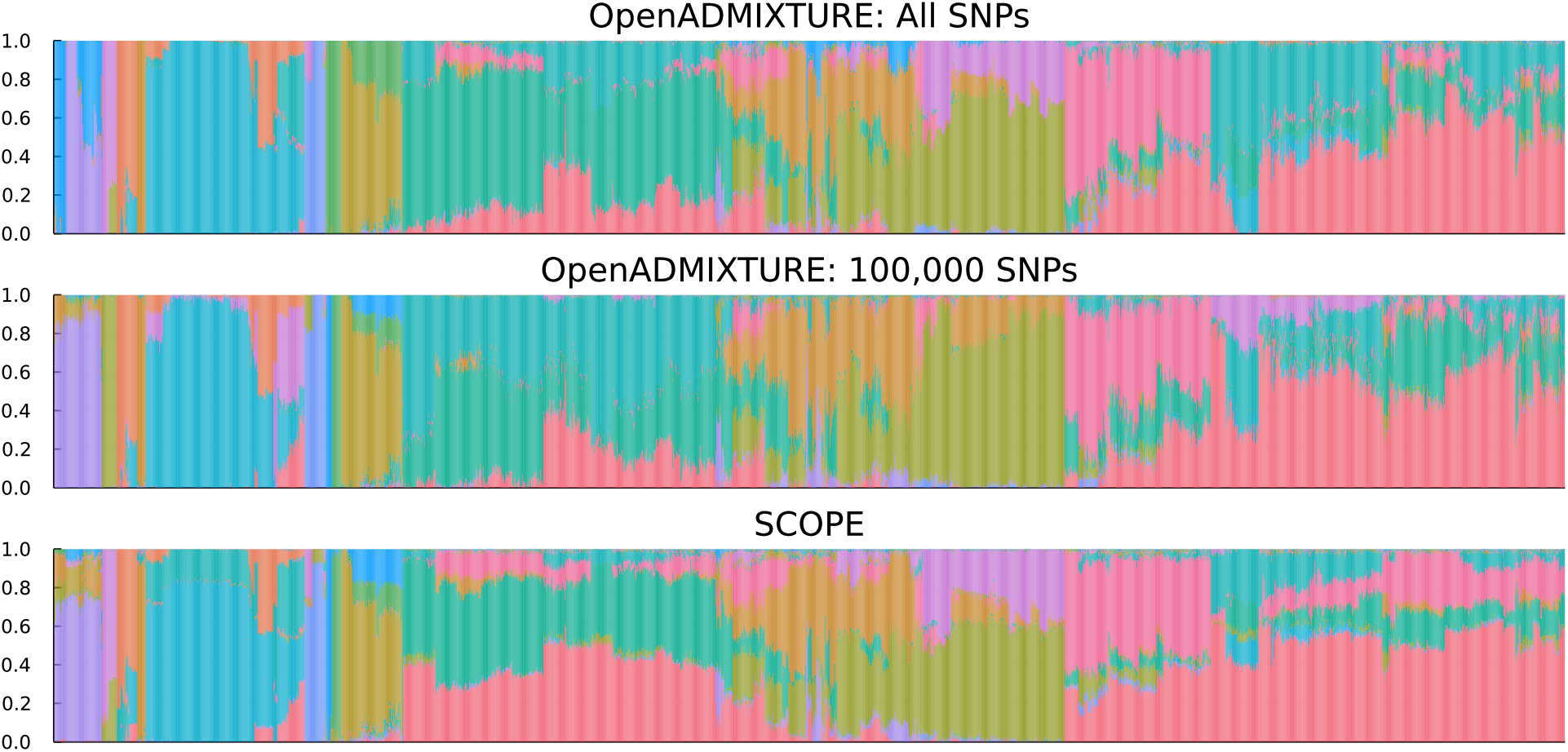
Estimated ancestry of HO data samples using (a) OpenADMIXTURE with all SNPs, (b) OpenADMIXTURE with 100,000 AIMs, and (c) SCOPE. These are stacked bar plots, with the y-axis indicating the proportion of total ancestry. The x-axis runs over all samples.

### 3.2 Computation Times and Maximum Memory Requirements

Most of our numerical experiments were run on Amazon Web Services (AWS). Table 9 lists the hardware instances invoked for computation. For our GPU experiments, we used two types of GPUs. The first, Nvidia A10G in a g5.4xlarge instance, is a moderate-grade GPU designed for low-cost performance. The second, Nvidia V100 in a p3.2xlarge instance, is specialized for scientific computing. The main difference between the two is double precision performance. By design, double precision performance on Nvidia A10G is 32 times slower than single precision, while double precision on Nvidia V100 is only twice as slow as single precision.

**Table 9:** Hardware instances used for experiments. The listed financial costs reflect on-demand instances in 2022 for the Oregon region, where all experiments were run.

| Instance | # of CPUs | Memory (GB) | GPU (Memory in GB) | Cost (\$/hr) |
| --- | --- | --- | --- | --- |
| g5.4xlarge | 16 | 64 | Nvidia A10G (24) | 1.624 |
| p3.2xlarge | 8 | 61 | Nvidia V100 (16) | 3.060 |
| c6i.16xlarge | 64 | 128 | -None- | 2.720 |
| r6i.16xlarge | 64 | 512 | -None- | 4.032 |

#### 3.2.1 Comparison to SCOPE

It is instructive to compare the runtime of OpenADMIXTURE (which pipelines SKFR followed by admixture estimation) to SCOPE (the previously most scalable method). For our pipeline on the TGP data, when all 16 available threads are used in a g5.4xlarge instance, a single SKFR run takes 1 minute 36 seconds. Filtering takes less than a minute. The subsequent run of admixture estimation takes less than 5 minutes with 100,000 or fewer SNPs. Cumulatively, the pipeline takes fewer than 7 minutes. On the other hand, a SCOPE run on the TGP data takes slightly over 16 minutes on the same hardware.

For the UKB data with *K* = 4, a single run of SKFR takes 44 minutes on a c6i.16xlarge instance with 128 GB memory. The maximum memory footprint is 73.2 GB. Creating the PLINK files containing only the selected AIMs takes less than 10 minutes. The subsequent admixture estimation run with 100,000 selected SNPs takes 29 minutes on a V100 GPU. Thus, the total pipeline runtime, invoked by a single OpenADMIXTURE call, was under 83 minutes. SCOPE took a similar 91 minutes to run on the UKB data set. Since SCOPE’s memory requirement for this data set is 250 GB, it had to be run on a more expensive r6i.16xlarge instance. In summary, total computation times are comparable, but OpenADMIXTURE is clearly less memory intensive than SCOPE.

#### 3.2.2 Runtime Improvement versus the Original ADMIXTURE Software

Although its based closely on the same methodology, OpenADMIXTURE delivers better performance than the original ADMIXTURE software. Table 10 records the per-iteration times of various admixture estimation routines on the TGP data with 100,000 AIMs. CPU and A10G GPU experiments were performed on a AWS g5.4xlarge hardware instance; V100 GPU experiments were performed on a AWS p3.2xlarge hardware instance. OpenADMIXTURE software, when restricted to CPUs, is 2.8 times faster on a single thread and 8 times faster in a 16-thread run compared to the original single-threaded ADMIXTURE. When a GPU is available, OpenADMIXTURE accelerates computation by another factor of 2 to 4, depending of course on the GPU hardware and the floating-point precision invoked.

**Table 10:** Per-iteration time for admixture estimation on a 100,000-SNP subset of the TGP data. CPU and A10G GPU experiments performed on a AWS g5.4xlarge hardware instance; V100 GPU experiments performed on a AWS p3.2xlarge hardware instance.

| Mode | Time (secs) |
| --- | --- |
| Single precision OpenADMIXTURE on V100 GPU | 1.4 |
| Double precision OpenADMIXTURE on V100 GPU | 1.5 |
| Single precision OpenADMIXTURE on A10G GPU | 1.6 |
| Double precision OpenADMIXTURE on A10G GPU | 2.8 |
| Double precision OpenADMIXTURE on CPU (16 threads) | 6.0 |
| Double precision OpenADMIXTURE on CPU (1 thread) | 28.5 |
| Double precision original ADMIXTURE on CPU (16 threads) | 48.0 |
| Double precision original ADMIXTURE on CPU (1 thread) | 79.0 |

**Table 11:**
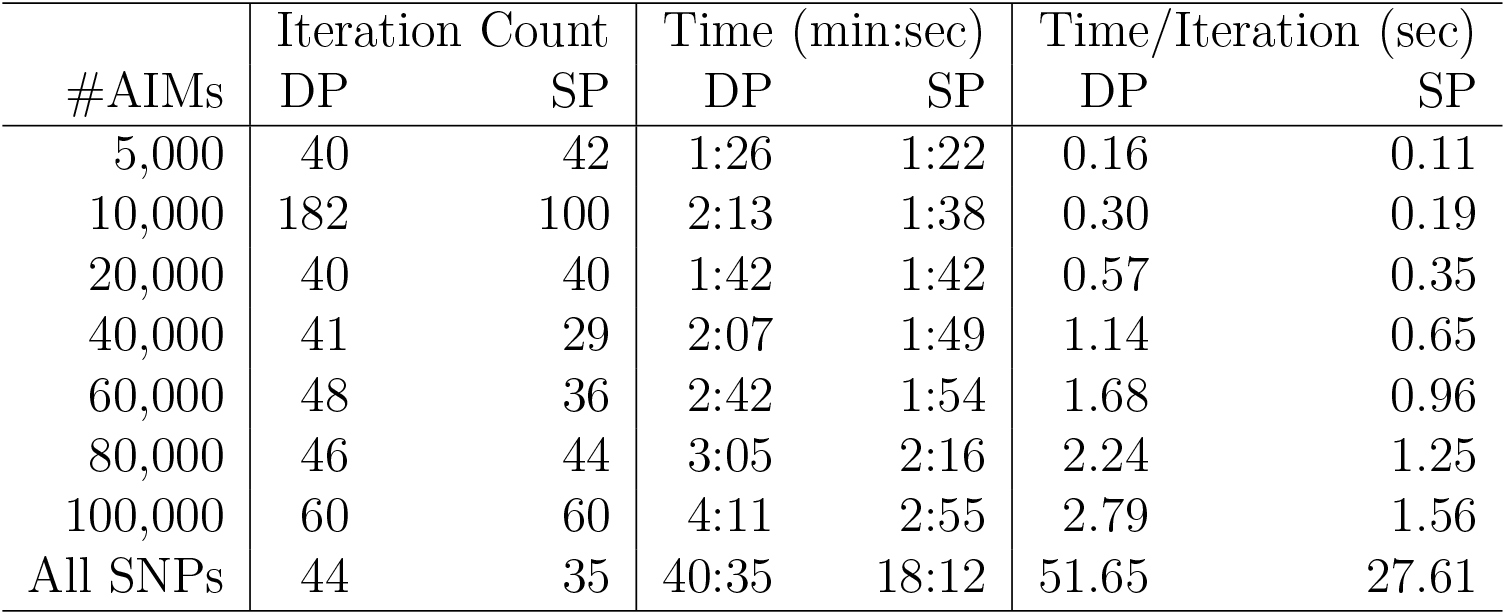
Comparison of iteration counts and runtimes for double precision (DP) versus single precision (SP) OpenADMIXTURE runs on the TGP data.

| #AIMs | Iteration Count |  | Time (min:sec) |  | Time/Iteration (sec) |  |
| --- | --- | --- | --- | --- | --- | --- |
|  | DP | SP | DP | SP | DP | SP |
| 5,000 | 40 | 42 | 1:26 | 1:22 | 0.16 | 0.11 |
| 10,000 | 182 | 100 | 2:13 | 1:38 | 0.30 | 0.19 |
| 20,000 | 40 | 40 | 1:42 | 1:42 | 0.57 | 0.35 |
| 40,000 | 41 | 29 | 2:07 | 1:49 | 1.14 | 0.65 |
| 60,000 | 48 | 36 | 2:42 | 1:54 | 1.68 | 0.96 |
| 80,000 | 46 | 44 | 3:05 | 2:16 | 2.24 | 1.25 |
| 100,000 | 60 | 60 | 4:11 | 2:55 | 2.79 | 1.56 |
| All SNPs | 44 | 35 | 40:35 | 18:12 | 51.65 | 27.61 |

#### 3.2.3 Double Precision versus Single Precision

One tactic for improving the computational speed of OpenADMIXTURE is to rely on single precision arithmetic in sequential quadratic programming. The gradient and Hessian are computed in single precision, but the loglikelihood is computed in double precision to determine convergence.

## 4 Discussion

This paper presents a biobank-scalable, unsupervised pipeline for AIM selection and estimation of admixture proportions for modern genetic data sets. We have implemented this entire procedure in our package OpenADMIXTURE. The SKFR (sparse *K*-means with a feature ranking) component of the pipeline is highly parallelized and effective in AIM selection. SKFR’s unsupervised clustering is insensitive to a small fraction of labeling errors and admixed samples. SKFR also delivers an explicit ranking of AIMs. Our experiments suggest that 100,000 informative SNPs deliver better clusters than full SNP sets. Uninformative SNPs simply constitute noise that slows clustering and obscures subpopulations.

The second component of the pipeline, admixture proportion estimation, is an opensource re-implementation in the Julia programming language of the previous package ADMIXTURE. The original ADMIXTURE^6^ is widely used, with over 5000 Google citations. OpenADMIXTURE is up to 8 times faster than the original ADMIXTURE on CPUs with multithreading and even faster on computers with GPUs. Total computation time is comparable to SCOPE, the only other method currently scalable to biobank data. We have shown that both OpenADMIXTURE and SCOPE can analyze a data set with 500,000 individuals and 600,000 SNPs in well under 2 hours. The paper^12^ introducing SCOPE showed that it could analyze this data set in about 24 hours. However, SCOPE is now more efficient than the original version, and computers are more powerful.

The memory demands of OpenADMIXTURE are exceptionally light due to its systematic exploitation for both storage and computation of PLINK’s binary format for genotype data. OpenADMIXTURE’s peak memory footprint is less than 120% of the size of the file used to input the genotype data, which is less than 30% of the memory needs of SCOPE. Specifically, to analyze the above biobank data set, SCOPE requires 250 GB of RAM, while OpenADMIXTURE needs under 75 GB. OpenADMIXTURE is also based on a likelihood model that incorporates basic population genetics concepts.

The computational complexity *O*(*IJK*^2^) of Hessian computation is a bottleneck for OpenADMIXTURE in dealing with *K >* 20 populations. Limiting analysis to a small number of AIMs reduces runtimes but does not eliminate the *K*^2^ dependence. If it is found desirable to tackle problems with large *K*, then gradient ascent might be helpful. Unfortunately, gradient ascent subject to constraints tends to be slow unless one can determine a nearly optimal step size. Line searches along the gradient direction require repeated loglikelihood evaluations and are expensive. We defer resolution of this issue to future research.

We have ignored the possible biological insights offered by the AIMs selected by SKFR. The genomic locations of AIMs, and their relation to the ancestral populations of the samples, warrant further research. Selecting the number of clusters *K* is another issue. Methods based on cross-validation require repeated runs of the pipeline and may be impractical on biobank-scale data. Because OpenADMIXTURE relies on a likelihood model, determination of *K* is possible based on the Akaike information criterion (AIC) or the Bayesian information criterion (BIC). Choosing *K* for SKFR may benefit from standard gap statistics ^52^. Alternatively, one can run SKFR with a variety of *K* values, then check AIC or BIC with the selected clusters and AIMs using OpenADMIXTURE. Again, this is a question for future research.

OpenADMIXTURE offers the option of inferring admixture proportions based on the population allele frequencies available in reference panels such as TGP. This approach fixes the allele frequencies ***P***, and only updates admixture proportions ***Q*** by sequential quadratic programming. As this problem is convex and fully separable in samples, it can be easily applied to sample collections ranging from small to biobank scale.

In summary, OpenADMIXTURE is a substantial upgrade of ADMIXTURE. Although the full panoply of options already available in ADMIXTURE has not yet been implemented, the ADMIXTURE community will surely welcome an open-source version that can be co-operatively developed further. The OpenMendel tools that OpenADMIXTURE already exploits provide a clear path to further improvement. We also expect Julia’s parallelization ecosystem to expand over time. We solicit the suggestions and assistance of committed users in the ADMIXTURE community in our efforts.

